# Enhancing Mucosal-Associated Invariant T (MAIT) cell function and expansion with human selective serum

**DOI:** 10.1101/2022.10.10.511368

**Authors:** Daniel Labuz, Jackson Cacioppo, Kelin Li, Jeffrey Aubé, Daniel T. Leung

## Abstract

Mucosal-associated invariant T (MAIT) cells are promising innate-like lymphocytes with potential for use in anti-tumor immunotherapy. Existing MAIT cell expansion protocols are associated with potentially decremental phenotypic changes, including increased frequency of CD4+ MAIT cells and higher inhibitory receptor expression. In this study, we compared the effect on expansion of human MAIT cells of a serum replacement, Physiologix XF SR (Phx), with traditional serum fetal bovine serum (FBS) for supplementing RPMI-1640 media. Using flow cytometry, we found that Phx supported a significantly higher proliferative capacity for MAIT cells and resulted in a lower frequency of CD4+ MAIT cells, which have been associated with reduced Th1 effector and cytolytic functions. We saw that culturing MAIT cells in Phx led to better survival of MAIT cells and lower frequency of PD-1+ MAIT cells compared to FBS-supplemented media. Functionally, we saw that Phx supplementation was associated with a higher frequency of IFN-γ+ MAIT cells after stimulation with *E. coli* compared to FBS-supplemented RPMI. In conclusion, we show that MAIT cells cultured in Phx have higher proliferative capacity, lower expression of inhibitory receptors, and have higher capacity to produce IFN-γ after *E. coli* stimulation, compared to FBS-supplemented RPMI. This work shows that expanding MAIT cells with Phx compared to FBS-supplemented RPMI result in a more functionally desirable MAIT cell for future anti-tumor immunotherapy.

## Introduction

T cells require specific nutrients and environment for optimal function and proliferation,^1,2^ and recent advances in our understanding of immunometabolism have uncovered cellular mechanisms that enable T cells for rapid cytokine production.^3^ These mechanisms include the switch from glycolytic and oxidative phosphorylation (OXPHOS) following T cell activation.^1,2,4^ In particular, glucose is a crucial nutrient for the efficient production of cytokines, and uptake of glucose is rapidly induced after T cell activation.^5,6^ For swift production of energy, the glucose transporter GLUT1 increases in expression in murine conventional CD4+ T cells following TCR stimulation.^7^ In addition, unconventional innate-like T cells, such as murine natural killer (NK) T cells also require glucose for optimal cytokine production.^8^

Mucosal-associated invariant T (MAIT) cells are an innate-like T cell that rapidly produces effector molecules such as interferon (IFN)-γ and granzyme B following activation. MAIT cells can be activated in a TCR-dependent manner through interaction with microbial derived ligand 5-OP-RU loaded on to major histocompatibility complex class I-related gene protein (MR1) or TCR-independent manner with interleukin (IL)-12 and IL-18. Previously, it was shown that MAIT cells have a lower metabolic rate than conventional T cells and show significantly higher glycolytic activity following TCR and cytokine stimulation compared to non-MAIT CD8+ effector memory cells.^9^ GLUT1 expression in MAIT cells increases following TCR and cytokine (IL-12 and IL-18) stimulation and 2NBDG expression increases following TCR stimulation but not cytokine stimulation.^9^ When glycolysis is disrupted with glucose analog 2DG, stimulated MAIT cells produced significantly lower IFN-γ but not tumor necrosis factor (TNF)-α.^9^ These studies suggest that cytokine production in MAIT cells may vary depending on the nutrient composition of their environment.

MAIT cells have the potential to be an HLA-independent, pan-cancer immunotherapy^10,11^, and recent work has examined methods for expanding MAIT cells while preserving cytolytic function.^12^ The majority of studies on expanded MAIT cells have used fetal bovine serum (FBS) as the major component of media supplementation, which is relatively inexpensive and effective for functional and proliferative experiments. While others have shown effective proliferation of MAIT cells using 5-A-RU with methylglyoxal (MeG), IL-2, IL-7, and the serum replacement CTS (Gibco).^12,13^ The goal of this study was to examine the expansion ability of MAIT cells in culture supplemented with a novel serum replacement reagent Physiologix (Phx, detailed below), previously shown to enhance CAR-T cell survival and function *in vivo*.^14^ We compared the proliferation, function, and exhaustion markers of MAIT cells cultured using Phx in comparison to FBS as a supplement to conventional RPMI media. We found that use of Phx was associated with greater proliferation capacity and increased IFN-γ production in MAIT cells following stimulation.

## Materials and Methods

### Human subjects and processing of peripheral blood mononuclear cells for assays

We obtained blood from discarded leukocyte filters and apheresis cones from healthy anonymous blood donors, and isolated PBMCs by density gradient centrifugation using Ficoll-paque PLUS (Cytiva). We froze isolated PBMCs <10×10^6^ cell/mL in a medium containing 30% FBS, 60% RPMI-1640 (Gibco), and 10% DMSO. We thawed 1 mL aliquots of frozen PBMCs rapidly in a 37 ºC water bath and centrifuged in 10 mL warm RPMI-1640 media. We washed PBMCs twice in warm RPMI-1640 before proceeding to assays comparing either FBS (Gibco) or Physiologix XF SR (Nucleus Biologics) supplemented RPMI-1640.

### Media culturing test

We thawed PBMCs from cryovials and washed as previously described above and seeded 2.5×10^6^ cells/mL in RPMI-1640 with 10% FBS or 2% Physiologix XF SR in 5 separate wells corresponding to time of incubation. We stained extracellularly with fixable viability dye eFluor™ 780 (eBioscience), anti-CD3-PerCP-Cy-5.5 (Biolegend), anti-CD8-BV605 (Biolegend), anti-CD4-BV510 (Biolegend), anti-Vα7.2-PE-Cy7 (Biolegend), anti-LAG-3-BV786 (Biolegend), anti-CD69-APC (Biolegend), anti-CD25-PE-Cy5 (BD Biosciences), anti-PD-1-Alexa-Fluor-700 (Biolegend), anti-CD161-BV605 (Biolegend), anti-CD69-PE-Cy5 (Biolegend), anti-TCRVδ2-PE-Texas-Red (Biolegend), anti-Fas-BV711 (Biolegend), and anti-human MR1-5-OP-RU Tetramer (NIH Tetramer Core Facility). We collected data using a 5-laser LSR-Fortessa flow cytometer (BD Biosciences) and the flow data was analyzed using FlowJo software v10 (Tree Star, Inc. Ashland, OR).

### *E. coli* stimulation

We thawed PBMCs from cryovials and washed as previously described above and seeded 2.5×10^6^ cells/mL in RPMI-1640 with 10% FBS or 2% Physiologix XF SR. We stimulated PBMCs with 10 multiplicity of infection (MOI) of strain 1100-2 fixed *E*. coli for 20 hours and blocked extracellular transport with brefeldin A for a total of 4 hours. We stained extracellular with fixable viability dye eFluor™ 780 (eBioscience), anti-CD3-BUV395 (BD Biosciences), anti-CD8-BV605 (Biolegend), anti-CD4-BV510 (Biolegend), anti-Vα7.2-PE-Cy7 (Biolegend), anti-LAG-3-BV786 (Biolegend), anti-CD25-BV650 (Biolegend), anti-PD-1-PerCP-Cy-5.5 (Biolegend), anti-CD161-BV605 (Biolegend), anti-CD69-PE-Cy5 (Invitrogen), anti-TCRVδ2-PE-Texas-Red (Biolegend), anti-CD161-APC (Biolegend), and anti-human MR1-5-OP-RU Tetramer (NIH Tetramer Core Facility). The PBMCs were fixed and permeabilized using Foxp3 / Transcription factor kit (eBioscience) and stained intracellularly with anti-Granzyme-B-Alexa-Fluor-700 (Biolegend), anti-TNF-α-efluor-450 (Invitrogen), anti-IFN-γ-FITC (Biolegend), anti-T-bet-BV711 (Biolegend). We collected data using a 5-laser LSR-Fortessa flow cytometer (BD Biosciences) and the flow data was analyzed using FlowJo software v10 (Tree Star, Inc. Ashland, OR).

### Expansion of MAIT cells

We thawed PBMCs from cryovials and washed as described above and seeded 1×10^6^ cells/mL in RPMI-1640 with 10% FBS or 2% Physiologix XF SR in a total of 1 mL. We used a method of MAIT cell expansion as previously described^13^ with the following modifications: instead of using base medium of Immunocult-XF T cell expansion medium (Stemcell Technologies) we used RPMI-1640 and substituted CTS serum (Invitrogen) with 10% FBS or 2% Physiologix XF SR and Normocin/Gentamicin with 1% penicillin/streptomycin. We further supplemented the media with 5 ng/mL recombinant human (rh) IL-2 (Peprotech) and 10 ng/mL rhIL-7. We stimulated proliferation of MAIT cells with 10 nM 5-OP-RU on Day 0, 5, and 10, and 50% of the media was exchanged every 2-3 days with fresh cytokines. On days 0, 7, and 14 100 µL was analyzed for MAIT cell count, frequency, activation, and inhibitory receptor markers. For extracellular staining we used: Zombie UV (Biolegend), anti-CD3-BUV395 (BD Biosciences), anti-CD8-BV605 (Biolegend), anti-CD4-BUV496 (BD Biosciences), anti-Vα7.2-BV711 (Biolegend), anti-LAG-3-BV786 (Biolegend), anti-CD69-BUV563 (BD Biosciences), anti-CD161-PE-Dazzle-594 (Biolegend), anti-TIM-3-BV421 (Biolegend), and anti-human MR1-5-OP-RU Tetramer (NIH Tetramer Core Facility). Sample data were acquired with a 5-laser Cytek Aurora flow cytometer (Cytek) and analyzed using FlowJo software v10 (Tree Star, Inc. Ashland, OR).

### Statistical analysis

For comparisons of FBS and Physiologix XF SR, a paired Wilcoxon Ranked Sums test was used. GraphPad Prism 8.3.0 software was used for statistical analysis of the flow cytometry data and *p* < 0.05 was considered statistically significant.

## Results

### Ligand-activated MAIT cells in Phx-supplemented RPMI had greater proliferative capacity and lower expression of inhibitory receptor TIM-3

We first examined the ability of Phx to proliferate MAIT cells *in vitro* using a published method^13^ of activating MAIT cells with the ligand 5-OP-RU three times over the course of 10 days (Figure S1A). We defined MAIT cells as live CD3+ Vα7.2+ MR1-Tetramer+ (Figure 1A). We supplemented RPMI-1640 with 2% Physiologix XF SR (Phx) or 10% FBS (FBS). We saw a significant increase in MAIT frequency and MAIT cell count in Phx compared to FBS-supplemented RPMI on day 7 (p = 0.0005) and day 14 (p = 0.0005) (Figure 1B-D). Next, we examined the proportion of CD4+ MAIT cells, given that CD4+ MAIT cells are known to have reduced Th1 effector and cytolytic functions.^15^ We saw significantly lower CD4+ MAIT cells in Phx compared to FBS-supplemented RPMI at day 7 and day 14 (Figure 1E). We also examined the expression of inhibitory receptors, which have been associated with reduced functionality among MAIT cells in colon tumors.^16^ We found that MAIT cells cultured in Phx has significantly less TIM-3+ MAIT cells after day 7 and day 14 compared to FBS-supplemented RPMI, though we saw no differences in LAG-3+ MAIT cells (Figure 1F-I). Inhibitory receptor expression often coincides with activation in MAIT cells^17^, and we see significantly lower CD69 expression in MAIT cells at day 14 in Phx compared to FBS-supplemented RPMI (Figure S1B-C). Overall, Phx supplementation resulted in greater proliferative capacity, a lower proportion of CD4+ MAIT cells, and lower frequency of TIM-3+ MAIT cells compared to FBS-supplemented RPMI.

**Figure 1.**
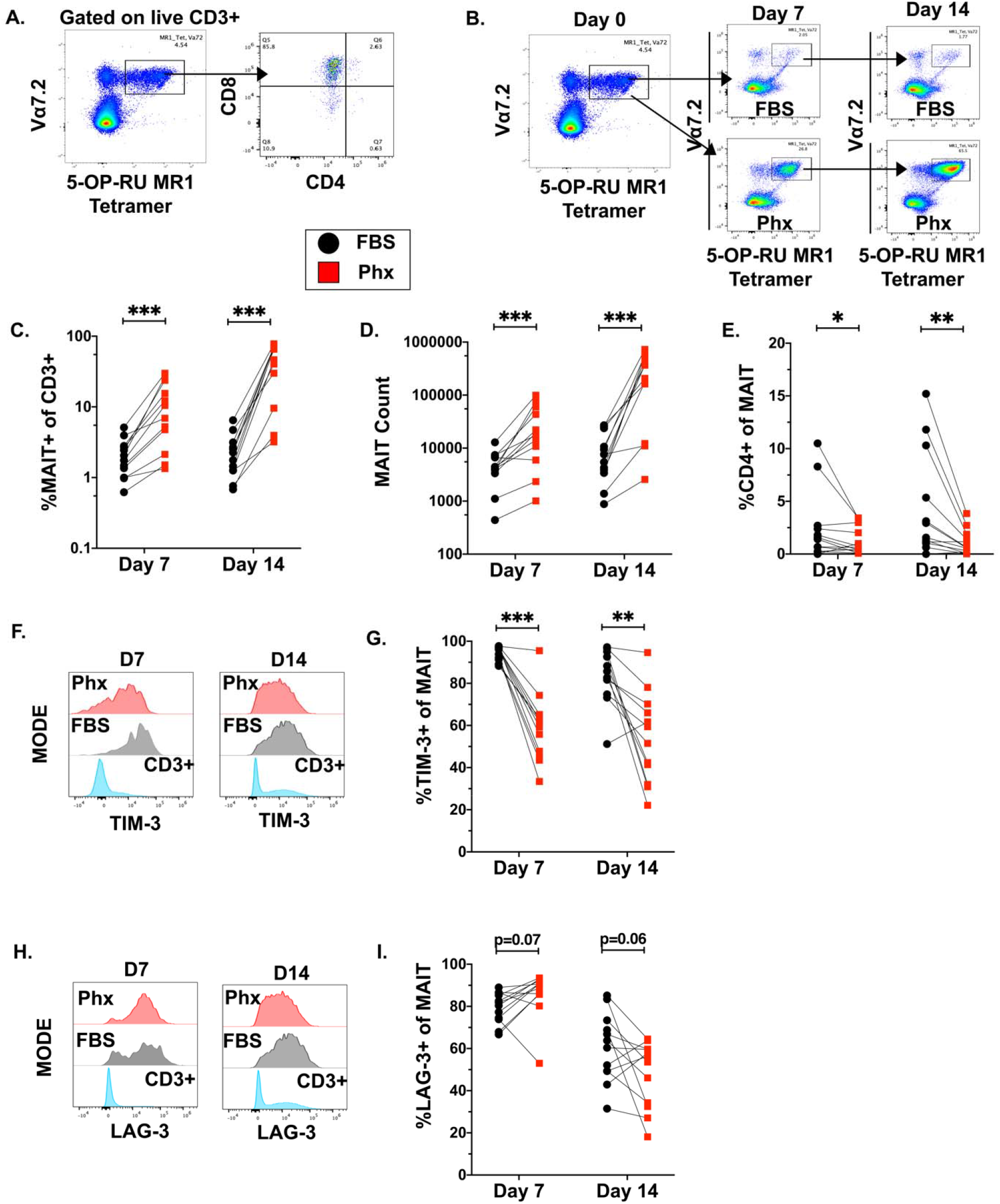
Phx increases proliferation capacity of MAIT cells and shows lower CD4+ MAIT cell frequency and lower expression of TIM-3 compared to FBS-supplemented RPMI. Gating strategy of MAIT cells from Live CD3+ and CD4+ and CD8+ gating strategy of MAIT cells (A). Representative MAIT cell proliferation flow plot of a sample culture in Phx or FBS-supplemented RPMI (B). Comparison of frequency of MAIT cells after 7 days of proliferation and 14 days of proliferation between Phx (Red box) or FBS- (Black circle) supplemented RPMI (C). Comparison of count of MAIT cells after 7 days of proliferation and 14 days of proliferation between Phx or FBS-supplemented RPMI (D). Comparison of the frequency of CD4+ MAIT cells after 7 days of proliferation and 14 days of proliferation between Phx or FBS-supplemented RPMI (E). Frequency of TIM-3+ MAIT cells after 7 days or 14 days between Phx or FBS-supplemented RPMI (F-G). Frequency of LAG-3+ MAIT cells after 7 days or 14 days between Phx or FBS-supplemented RPMI (H-I). These data are reflective of 2 independent experiments with n=6 each. *-P<0.05, **-P<0.01, ***-P<0.001.

### Non-activated MAIT cells cultured in Phx-supplemented RPMI had better persistence and lower inhibitory receptor expression

To determine the impact of Phx on conventional and unconventional T cells survival, we cultured primary T cells from 4 blood donors for 8 days. We do not see any significant differences in CD3+ frequency or total CD3+ cells between Phx and FBS-supplemented RPMI at any timepoint measured (Figure 2A-C). However, when we examined the impact on MAIT cells, we saw significantly higher frequency and count of MAIT cells at all timepoints in Phx compared to FBS-supplemented RPMI (Figure 2D-E). On the other hand, we did not see any significant differences in γδT Vδ2 count or frequency between Phx and FBS-supplemented RPMI at any timepoint (Figure S2A-B). We then looked at expression of inhibitory receptor PD-1 as it has been implicated in exhausted MAIT cells in tumors.^18^ We saw that there was significantly lower frequency of PD-1+ MAIT cells in the Phx at all timepoints (Figure 2F-G). Similarly, we saw significantly lower PD-1+ γδT Vδ2 cells in Phx compared to FBS-supplemented RPMI (Figure S2C-D). Lastly, we saw significantly lower LAG-3+ and TIM-3+ expression on MAIT cells in Phx compared to FBS-supplemented RPMI (Figure 2H-I). Overall, we saw no differences in frequency or counts of conventional T cells or γδT Vδ2 cells between Phx or FBS-supplemented RPMI, but noted significantly higher frequency and count of MAIT cells with lower expression of inhibitory receptors in Phx compared to FBS-supplemented RPMI.

**Figure 2.**
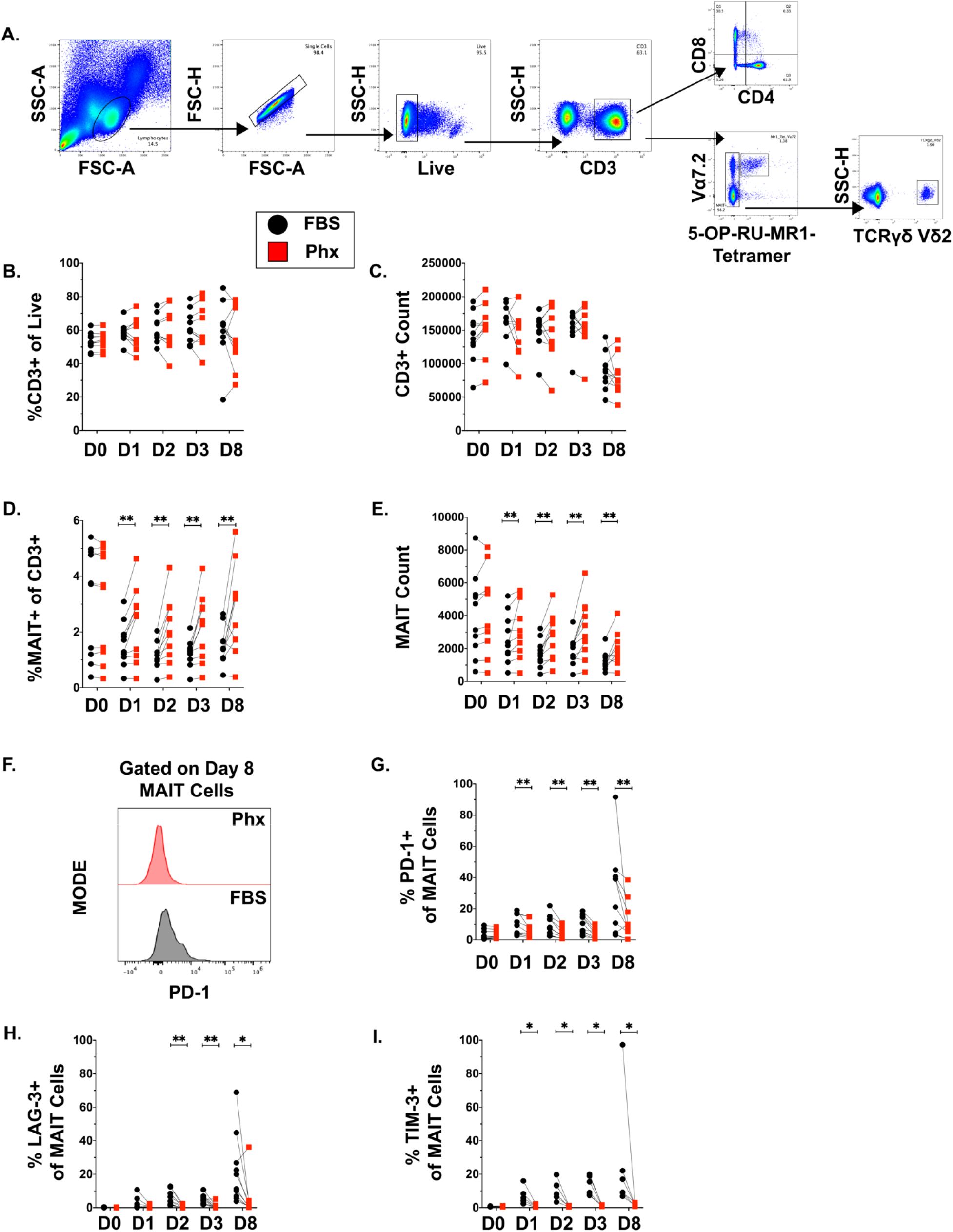
Phx-supplementation shows lower MAIT cell PD-1 expression following eight days culture compared to FBS-supplementation. Gating strategy of CD3+, CD4+, CD8+, MAIT, and TCRVδ2 T cells (A). Comparison of CD3+ frequency of live cells and CD3+ counts of 4 time-points: day 1, day 2, day 3, and day 8 between Phx (Red box) and FBS- (Black circle) supplemented RPMI (B&C). Comparison of MAIT cell frequency of CD3+ T cells and MAIT cell counts across 4 timepoints between Phx and FBS-supplemented RPMI (D&E). Frequency of PD-1+ MAIT cells after 8 days of culturing in Phx or FBS-supplemented RPMI (F&G). These data are reflective of 2 experiments with n=5. *-P<0.05, **-P<0.01, ***-P<0.001.

### *E. coli*-stimulated MAIT cells in Phx-supplemented RPMI had greater IFN-_γ_ expression compared to FBS-supplemented RPMI

We next looked at the effect of Phx on the effector molecule production of MAIT cells following stimulation with *E. coli*, a relevant microbe with capacity to produce the MAIT-stimulating ligand We saw no changes in MAIT cell frequency after stimulation with *E. coli* between Phx and FBS-supplemented RPMI (Figure 3A). We looked at activation markers and saw no differences in frequency of CD69+ and CD25+ MAIT cells between Phx and FBS-supplemented RPMI following *E. coli* stimulation (Figure 3B-C). We also did not see any significant differences in frequency of LAG-3+ or PD-1+ MAIT cells, but lower frequency of TIM-3+ MAIT cells (Figure 3D-F). Notably, we saw significantly higher IFN-γ expression following *E. coli* stimulation in Phx compared to FBS-supplemented RPMI (Figure 3G-H) but no significant differences in TNF-α or Granzyme B expression (Figure 3I-J), nor in the expression of T-bet which is implicated in the regulation of IFN-γ expression (Figure 3K). Taken together, Phx supplementation of *E. coli* stimulated MAIT cells resulted in lower frequency of TIM-3+ MAIT cells and higher IFN-γ expression compared to FBS supplementation, but no differences seen in activation markers, LAG-3 and PD-1, or T-bet expression.

**Figure 3.**
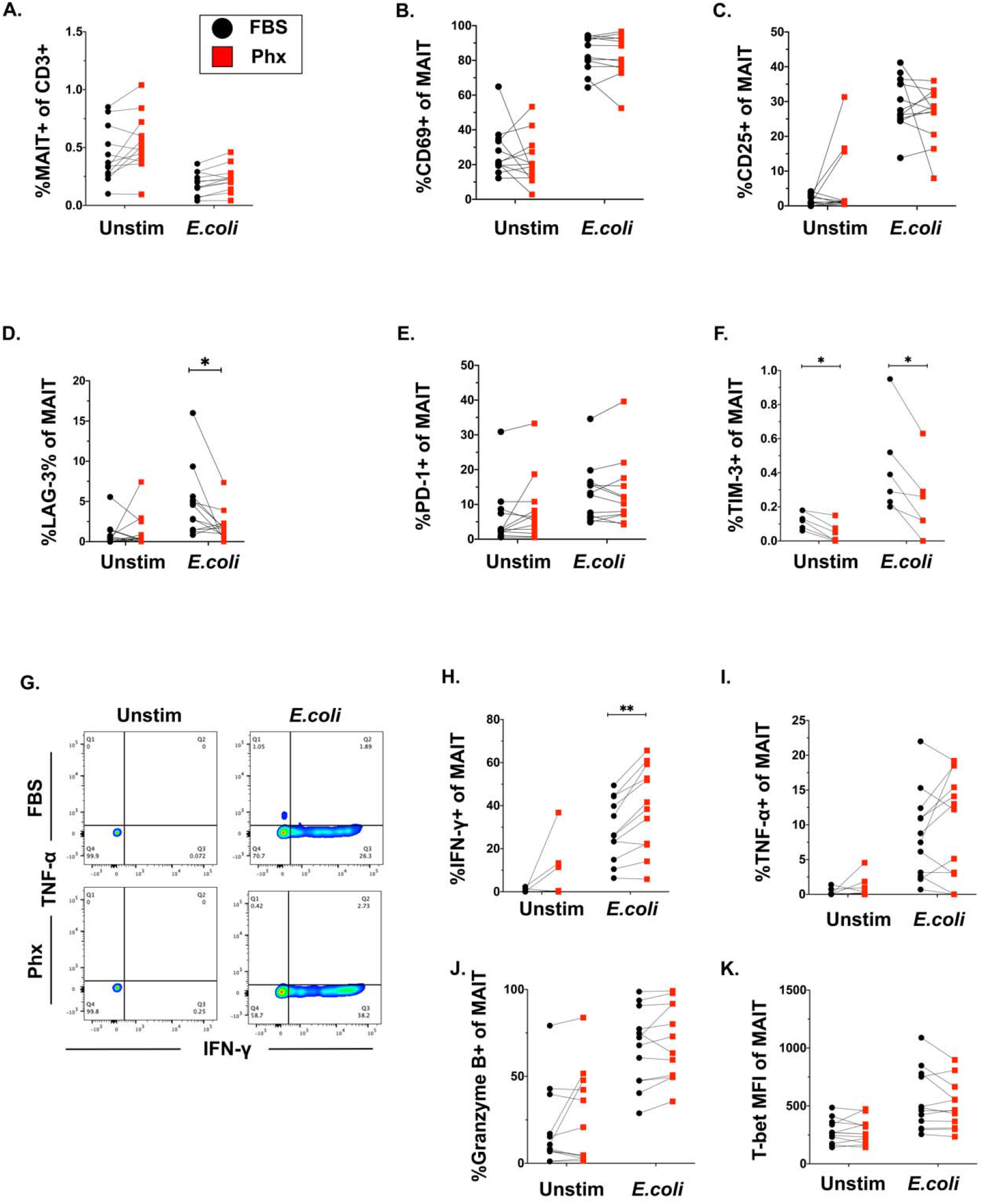
Phx shows higher IFN-_γ_ production in MAIT cells after *E. coli* stimulation compared to FBS-supplemented RPMI. Frequency of MAIT cells after 20 hours with or without 10 MOI *E. coli* stimulation in Phx or FBS-supplemented RPMI (A). Comparison of frequency of CD69+, CD25+, LAG-3+, PD-1+, and TIM-3+ MAIT cells after 20 hours with or without 10 MOI *E. coli* stimulation in Phx or FBS-supplemented RPMI (B-F). Gating strategy and frequency of TNF-α+ and IFN-γ+ MAIT cells following 20 hours with or without 10 MOI *E. coli* stimulation in Phx or FBS-supplemented RPMI (G-I). Frequency of Granzyme-B+ MAIT cells after 20 hours with or without 10 MOI *E. coli* stimulation in Phx or FBS-supplemented RPMI (J). Comparison of MFI of T-bet expression in MAIT cells after 20 hours with or without 10 MOI *E. coli* stimulation in Phx or FBS-supplemented RPMI (K). These data are reflective of 2 independent experiments with n=6 each. *-P<0.05, **-P<0.01, ***-P<0.001.

## Discussion

In this study, we present a method to expand MAIT cells isolated from peripheral blood utilizing a novel serum substitute. We show that compared to FBS, Phx supplementation results in greater expansion, and a higher cytokine potential following *E. coli* stimulation, while maintaining a more biologically similar phenotype with lower expression of inhibitory receptor expression in expanded MAIT cells.

The potential use of expanded MAIT cells for anti-tumor immunotherapy has been suggested after recent studies demonstrating MR1 to be an important candidate target for human cancers.^10,19^ and demonstration of anti-tumor effects in various mouse models of MAIT immunotherapy.^20,21^ In humans, there is an increase in frequency of circulating CD4+ MAIT cells and a subsequent decrease of CD8+ MAIT cells compared to healthy controls suggesting that CD8+ T cells are infiltrating tumor sites.^22^ Our MAIT cell expansion protocol was able to mitigate the increase of frequency of CD4+ MAIT cells, thus possibly offering improved anti-tumor activity over previously-published protocols for MAIT expansion, which resulted in an increased frequency of CD4+ MAIT cells.^12^ The ability to efficiently expand MAIT cells *ex vivo* whilst maintaining a functional phenotype may support the development of new MAIT-based tumor immunotherapies.

The addition of a serum substitute such as Phx provides differential nutrients for MAIT cells in culture compared with FBS, and was associated with decreased expression of inhibitory receptors and an increased capacity for IFN-γ production after *E. coli* stimulation but not cytokine stimulation. Previous studies have shown that MAIT cell production of granzyme B and IFN-γ is affected by glucose availability, most strikingly following a combination of TCR and cytokine stimulation suggesting that the combination of both pathways is required.^9,23^ While the exact composition of Phx is unknown, studies using mass spectrometry showed Phx showed higher levels of metabolites such as glucose-1-phosphate and pyruvate compared to human serum, both of which can be used to generate ATP through the TCA cycle for oxidative phosphorylation purposes.^2,14^

Inhibitory receptors such as PD-1 and TIM-3 are part of the subset of receptors that are associated with T cell exhaustion and decreased effector function^24^, as seen with MAIT cells in hepatic carcinoma,^18^ and colorectal cancer.^16^ Our data shows that Phx is associated with lower TIM-3 expression during ligand-activated MAIT cell proliferation and lower PD-1, TIM-3, and LAG-3 in unstimulated MAIT cell culture. These data suggest that expanding and culturing MAIT cells with Phx-supplemented RPMI may results in lower levels of MAIT cell exhaustion, which is desirable for anti-tumor immunotherapy.

In conclusion, we show that use of Phx-supplemented RPMI results in a higher proliferation and IFN-γ production from MAIT cells compared to FBS-supplemented RPMI. In addition, we saw that Phx was associated with decreased PD-1, TIM-3, and LAG-3 in MAIT cell culture, and lower TIM-3 expression in MAIT cell proliferation, compared to FBS-supplemented RPMI. Our findings have the potential to inform the development strategies for *ex vivo* proliferation and maintenance of a functional phenotype of MAIT cells that could be used in future immunotherapies against cancers.

## Acknowledgements

This research was supported by the National Institutes of Health (AI130378 to D.T.L. and TL1TR002540 to D.L). We would like to thank Associated Regional and University Pathologists for the blood samples. We would also like to thank the staff of the University of Utah Flow Cytometry Core. We thank Nucleus Biologics for providing some of the Physiologix XF SR used in this study.

