## Supplemental Figures for "Enhancing Mucosal-Associated Invariant T (MAIT) cell function and expansion with human selective serum"

### Figure Legends

**Figure 1S. Phx lowers CD69 expression of proliferating MAIT cells compared to FBS-supplemented RPMI.** Experimental layout of MAIT cell expansion protocol (A). Representative histogram of CD69 expression of Day 14 proliferated MAIT (B). Comparison of CD69 expression of MAIT cells after 7 days of proliferation and 14 days of proliferation between Phx (Red box) or FBS- (Black circle) supplemented RPMI (C). These data are reflective of 2 independent experiments with n=6 each. *-P<0.05, **-P<0.01, ***-P<0.001.

**Figure S2. Phx-supplementation does not impact TCRVδ2 T cells persistence but decreases PD-1 expression in culture compared to FBS-supplementation.** Comparison of TCRVδ2 T cells frequency of live cells and TCRVδ2 T cells counts of 4 time-points: day 1, day 2, day 3, and day 8 between Phx (Red box) and FBS- (Black circle) supplemented RPMI (A&B). Frequency of PD1+ TCRVδ2 T cells after 8 days of culturing in Phx or FBS-supplemented RPMI (C&D). These data are reflective of 2 experiments with n=5. *-P<0.05.

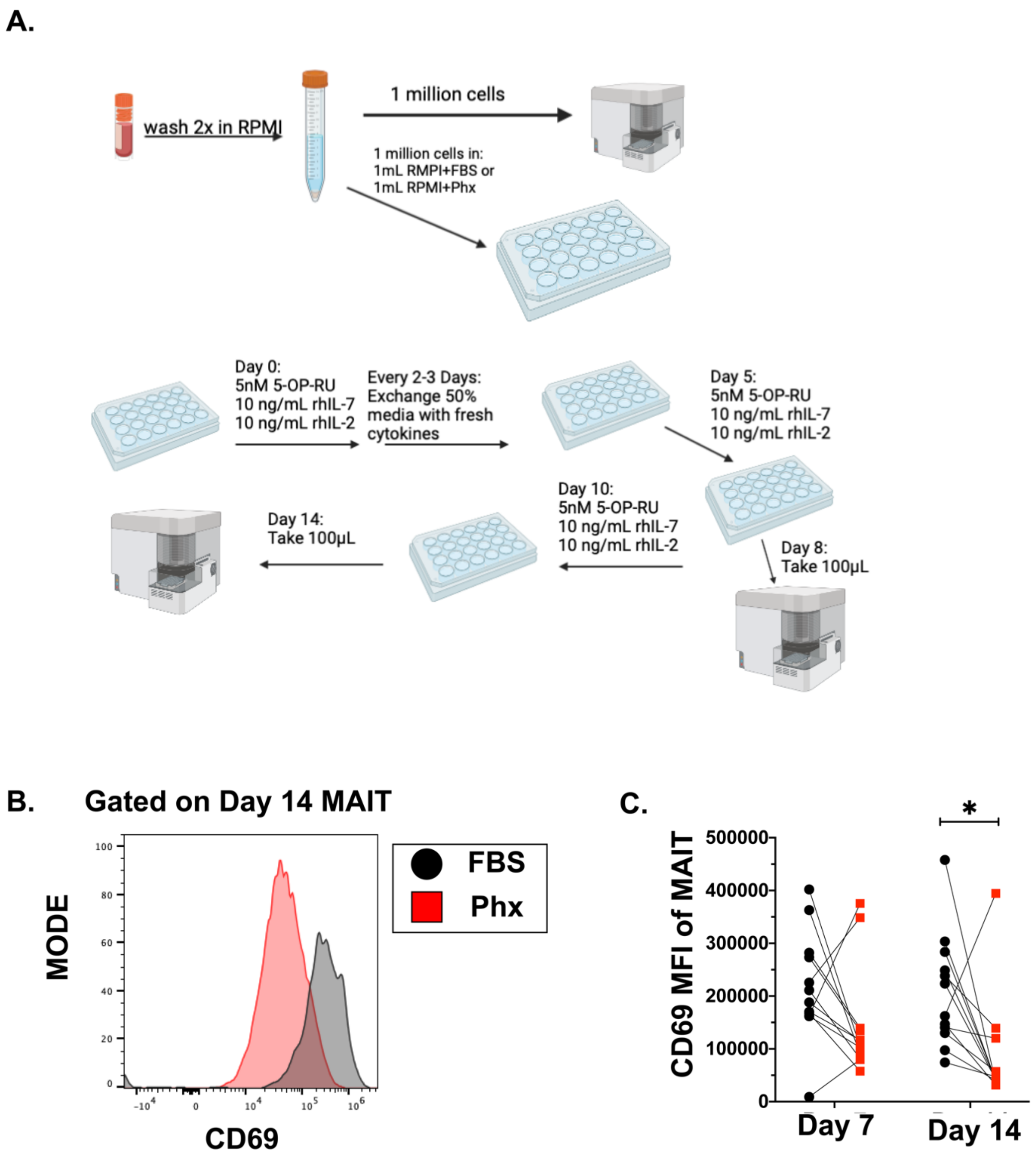
A.
wash 2x in RPMI
1 million cells
1 million cells in:
1mL RMPI+FBS or
1mL RPMI+Phx
Day 0:
5nM 5-OP-RU
10 ng/mL rhIL-7
10 ng/mL rhIL-2
Every 2-3 Days:
Exchange 50%
media with fresh
cytokines
Day 5:
5nM 5-OP-RU
10 ng/mL rhIL-7
10 ng/mL rhIL-2
Day 10:
5nM 5-OP-RU
10 ng/mL rhIL-7
10 ng/mL rhIL-2
Day 8:
Take 100μL
Day 14:
Take 100μL
B.
Gated on Day 14 MAIT
MODE
CD69
FBS
Phx
C.
CD69 MFI of MAIT
Day 7
Day 14
*

Figure S1.

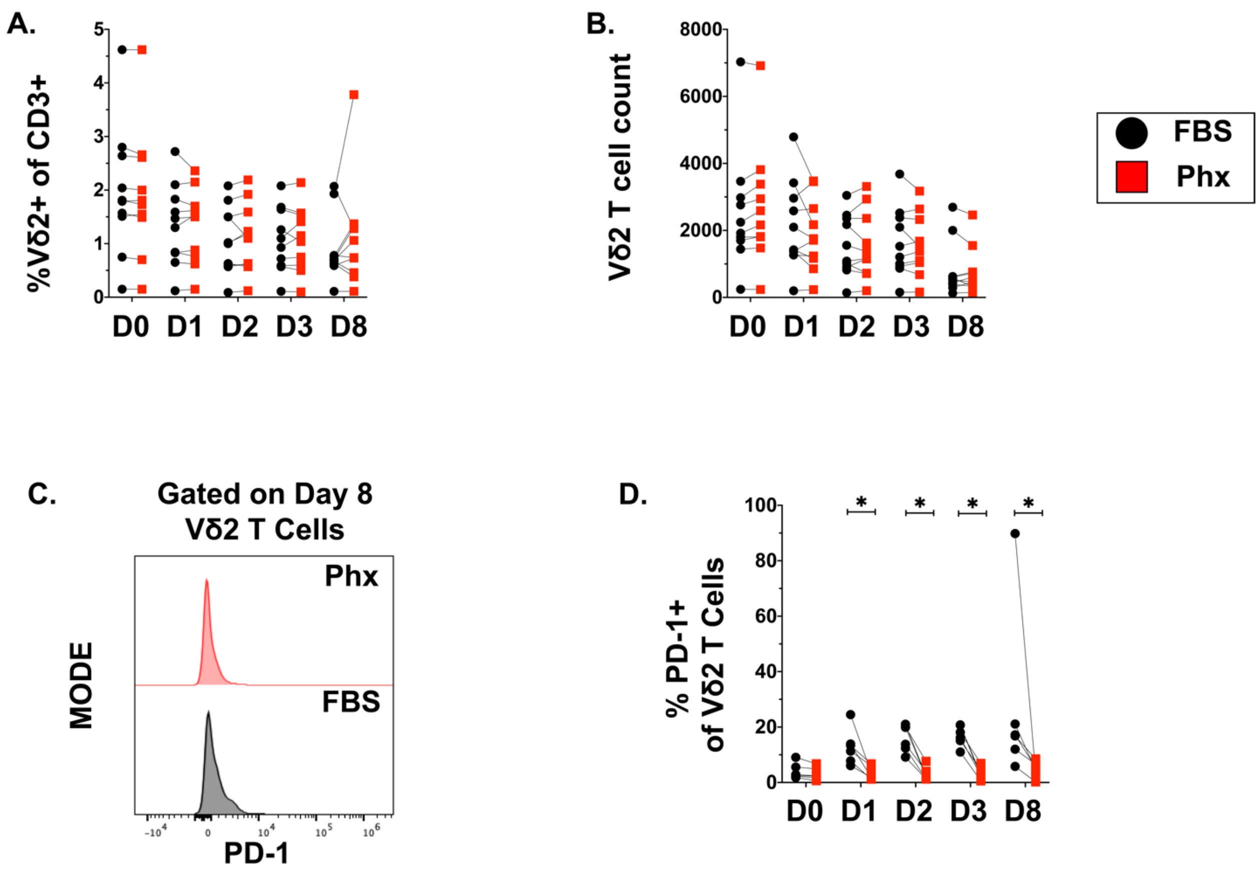
A.
%Vδ2+ of CD3+
5
4
3
2
1
0
D0 D1 D2 D3 D8
B.
Vδ2 T cell count
8000
6000
4000
2000
0
D0 D1 D2 D3 D8
FBS
Phx
C.
Gated on Day 8
Vδ2 T Cells
MODE
Phx
FBS
PD-1
D.
% PD-1+
of Vδ2 T Cells
100
80
60
40
20
0
*
*
*
*
D0 D1 D2 D3 D8

Figure S2.
